# Intrinsic cortical dynamics dominate population responses to natural images across human visual cortex

**DOI:** 10.1101/008961

**Authors:** Linda Henriksson, Seyed-Mahdi Khaligh-Razavi, Kendrick Kay, Nikolaus Kriegeskorte

## Abstract

Intrinsic cortical dynamics are thought to underlie trial-to-trial variability of visually evoked responses in animal models. Understanding their function in the context of sensory processing and representation is a major current challenge. Here we report that intrinsic cortical dynamics strongly affect the representational geometry of a brain region, as reflected in response-pattern dissimilarities, and exaggerate the similarity of representations between brain regions. We characterized the representations in several human visual areas by representational dissimilarity matrices (RDMs) constructed from fMRI response-patterns for natural image stimuli. The RDMs of different visual areas were highly similar when the response-patterns were estimated on the basis of the same trials (sharing intrinsic cortical dynamics), and quite distinct when patterns were estimated on the basis of separate trials (sharing only the stimulus-driven component). We show that the greater similarity of the representational geometries can be explained by the coherent fluctuations of regional-mean activation within visual cortex, reflecting intrinsic dynamics. Using separate trials to study stimulus-driven representations revealed clearer distinctions between the representational geometries: a Gabor wavelet pyramid model explained representational geometry in visual areas V1–3 and a categorical animate– inanimate model in the object-responsive lateral occipital cortex.

## INTRODUCTION

Visual stimulation has been shown in animal models to only slightly modulate ongoing cortical dynamics in the visual cortex (Arieli et al. 1996; Fiser et al. 2004). Trial-to-trial variability of evoked fMRI responses in human cortex has also been related to coherent intrinsic fluctuations of activity (Fox et al. 2006; Becker et al. 2011). However, the effect of intrinsic dynamics on visual representations and their functional role is not well understood.

The representational content of neuronal population codes is increasingly being investigated with pattern-information techniques (Kriegeskorte and Kreiman 2011). In this approach, stimulus-related activity patterns are interpreted as distributed representations of the stimuli. Functional magnetic resonance imaging (fMRI) enables us to image many areas simultaneously and to investigate the transformation of the representational space across stages of processing. However, previous studies have ignored the effect of intrinsic dynamics on comparisons of representations between different areas. That is, visual areas show prominent coherent fluctuations in spontaneous activity between the areas in the absence of visual stimulation (Shmuel and Leopold 2008), a phenomenon typically referred to in human brain imaging as functional connectivity (Nir et al. 2006; Fox and Raichle 2007). Resting-state functional connectivity between brain regions has also been shown to be highly similar to connectivity estimated based on fMRI response-pattern dissimilarities during task-performance (Ritchey et al. 2014). This has been interpreted as evidence for sub-networks of brain regions contributing to specific tasks. Resting-state functional connectivity measures and stimulus-related response-pattern dissimilarities may, however, have a common underlying component that has remained unrecognized. Here we show that estimates of stimulus-related response-pattern dissimilarities can be strongly affected by ongoing cortical dynamics.

The representational geometry of a visual area can be characterized by a representational dissimilarity matrix (RDM), which contains a representational distance for each pair of stimulus-related fMRI response patterns (Kriegeskorte, Mur and Bandettini 2008; Kriegeskorte and Kievit 2013). Representations in two brain regions can be compared by computing the correlation between their RDMs (Kriegeskorte 2009; Nili et al. 2014). Likewise, a brain RDM can be directly compared to a model RDM that captures the response-pattern dissimilarities in the internal representation of a computational model that processes the stimuli (Kriegeskorte 2009). Comparing visual representations between brain areas and processing stages in computational models can help us better understand the functional organization of the human visual cortex. The goal is to understand how the representational space is transformed across stages of processing. However, as shown here, coherent fluctuations of overall activation between two regions can make the apparent representational geometries of visual areas much more similar than the true underlying visual representations.

Figure 1 shows simulation results on the effect of coherent intrinsic response fluctuations on RDMs. In this simple simulation, primary visual cortex (V1) responded equally to three categories of stimuli (Bodies, Faces, Objects) whereas face-responsive fusiform face area (FFA) showed preference for Faces. The difference between the response profiles is reflected in their RDMs shown in Figure 1A. In Figure 1B, a coherent fluctuation component was added to the responses. The underlying stimulus-driven pattern variances remained the same but the visual areas now shared the fluctuation in the overall responsiveness across time. As a result, the RDMs of the two visual areas are highly similar (Fig. 1B), thus challenging the interpretation of the differences between the stimulus representations and highlighting the contribution of shared intrinsic response fluctuations on representational geometries.

**Figure 1.**
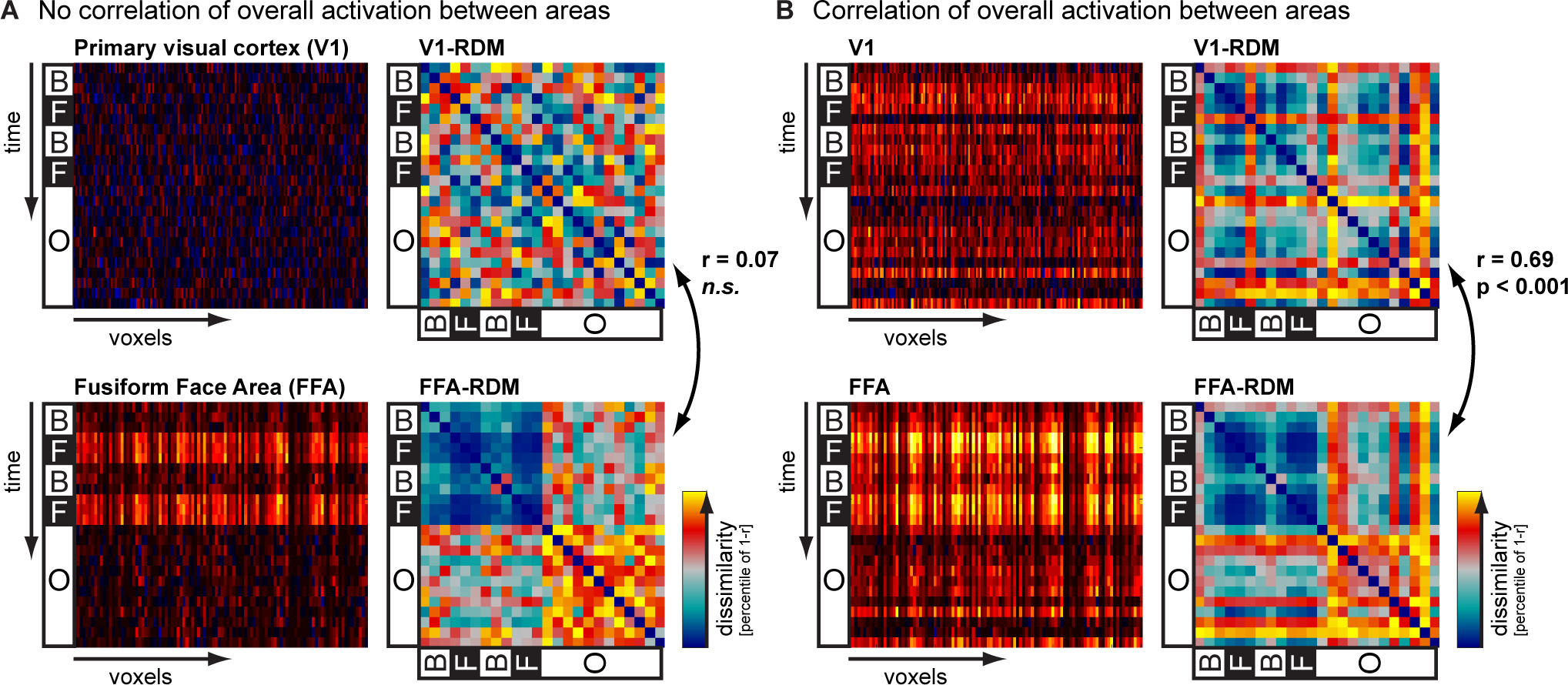
Simulation on the effects of coherent response fluctuations on RDM similarity. A) Simulated primary visual cortex (V1; 150 voxels) responds equally strongly to three categories of stimuli (B = Bodies, F = Faces, O = objects). The representational similarity matrix (RDM) captures the pair-wise representational distance between the response patterns for each stimulus with the V1 RDM showing no interesting structure. The simulated fusiform face area (FFA; 100 voxels) shows preference for Face-stimuli (F) and responds slightly more strongly also to the Bodies (B) than to Objects (O). This is reflected in the FFA RDM showing most similar response-patterns for the Faces. The V1 RDM and the FFA RDM are clearly different (Spearman’s rank correlation r = 0.07, not significant; condition-label randomization test (Kriegeskorte, Mur and Bandettini 2008)). B) A coherent response fluctuation component was added to V1 and FFA responses. The stimulus-driven patterns remained the same but the visual areas now shared the fluctuation in the overall responsiveness across time. As a result, the RDMs of the two visual areas are highly similar (Spearman’s rank correlation r = 0.69, p < 0.001; condition-label randomization test (Kriegeskorte, Mur and Bandettini 2008)). This shows that coherent response-pattern fluctuations can have a significant effect on visual representations as reflected in response-pattern dissimilarities.

In this study, we explored fMRI responses in human visual cortex to a large set (1750) of natural images (Kay et al. 2008). Figure 2 shows results illustrating coherent response fluctuations between visual areas in this data. Areas closer in cortex, and in the visual hierarchy, tended to exhibit greater functional connectivity. These fluctuations were unrelated to the stimuli and thus likely reflected intrinsic cortical dynamics. We will show that the coherent fluctuations have a strong effect on the representational similarity between visual areas, and that the true stimulus-driven component can be revealed by comparing RDMs constructed from response-patterns estimated on the basis of separate trials.

**Figure 2.**
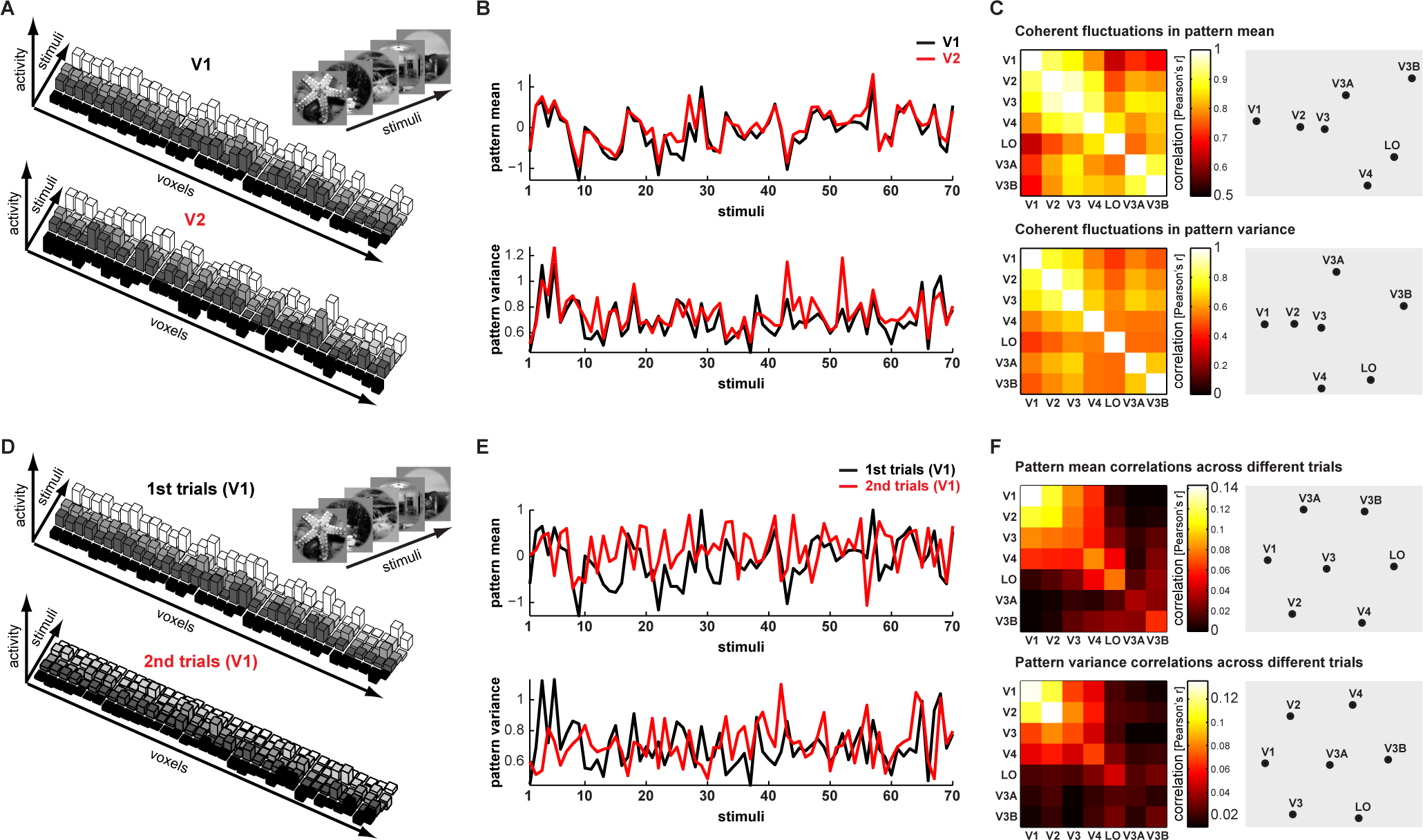
Coherent response-pattern fluctuations in natural image data. A) The response amplitudes for 5 natural images are shown for a subset of voxels in visual areas V1 and V2 of subject S1. The visual areas showed coherent dynamics in their response patterns. That is, the responses in all voxels in both V1 and V2 were low for the first stimulus, stronger for the second stimulus and again lower for the fourth stimulus. B) The mean and variance of the response pattern amplitudes for 70 natural stimuli are shown for visual areas V1 and V2. Both showed highly coherent dynamics between the visual areas. C) The matrices show the mean correlations between response pattern means (top row) and variances (bottom row) between all pair-wise comparisons of the visual areas. The matrices are also visualized using multidimensional-scaling arrangement. What emerged from the coherence of the response-pattern fluctuations is the hierarchy of visual areas. D–F) When the comparisons were done between repeated presentations of the same stimuli (separate trials), the correlations in the response-pattern-mean and variance were much lower. This suggests a significant contribution of the coherent response fluctuations on the similarity of RDMs from different visual areas in this data.

## MATERIALS AND METHODS

### Visual stimuli and fMRI data

The current study used fMRI data from a previously published study; for details on the visual stimuli, data acquisition and data pre-processing, please see Kay et al. (Kay *et al.* 2008) and Naselaris et al. (Naselaris et al. 2009). This data had been used as training data for voxel-receptive-field modelling. In short, the stimuli were 1750 gray-scale natural photographs that were masked with a 20°-diameter circle (for an example, see Figure 1A) and were presented for 1 s (flashed three times ON (200 ms)–OFF (200ms)–ON (200 ms)–OFF (200 ms)–ON (200ms)) with a 3-s fixation-only period between successive photographs and every eight trial being a null trial. Data from three subjects were analyzed (S1–S3). For each subject, the data had been collected in five separate scanning sessions with five experimental runs in each. Each experimental run consisted of 70 different natural images, each presented two times.

The data were pre-processed using an updated protocol which included slice-timing correction, motion correction, upsampling to (1.5 mm)^3 resolution and improved co-registration between the functional data sets. The data were modelled with a variant of the general linear model including discrete cosine basis set for the hemodynamic response function (HRF) estimation. Low-frequency noise fluctuations were accounted for by polynomials (for details, please see (Kay *et al.* 2008)). The beta weights characterizing the amplitude of the BOLD response to each stimulus were transformed to Z scores.

The regions-of-interests V1, V2, V3, V4, LO, V3A and V3B were based on independent localizer data based on retinotopic criteria (Naselaris *et al.* 2009). The analysis was restricted to voxels with signal-to-noise ratio greater than 1.5 (median value observed across all images). The regions-of-interest contained no overlapping voxels.

### Representational similarity analysis

The fMRI response patterns evoked by the different natural images were compared to each other using correlation distance, and all pairwise comparisons were assembled in a representational dissimilarity matrix (RDM) for each region (Kriegeskorte, Mur and Bandettini 2008; Nili *et al.* 2014). Figure 3B shows an example RDM. The RDMs were calculated separately for each experimental run with 70 different natural images in each. The measure for dissimilarity was correlation distance (1-Pearson linear correlation) between the response patterns.

**Figure 3.**
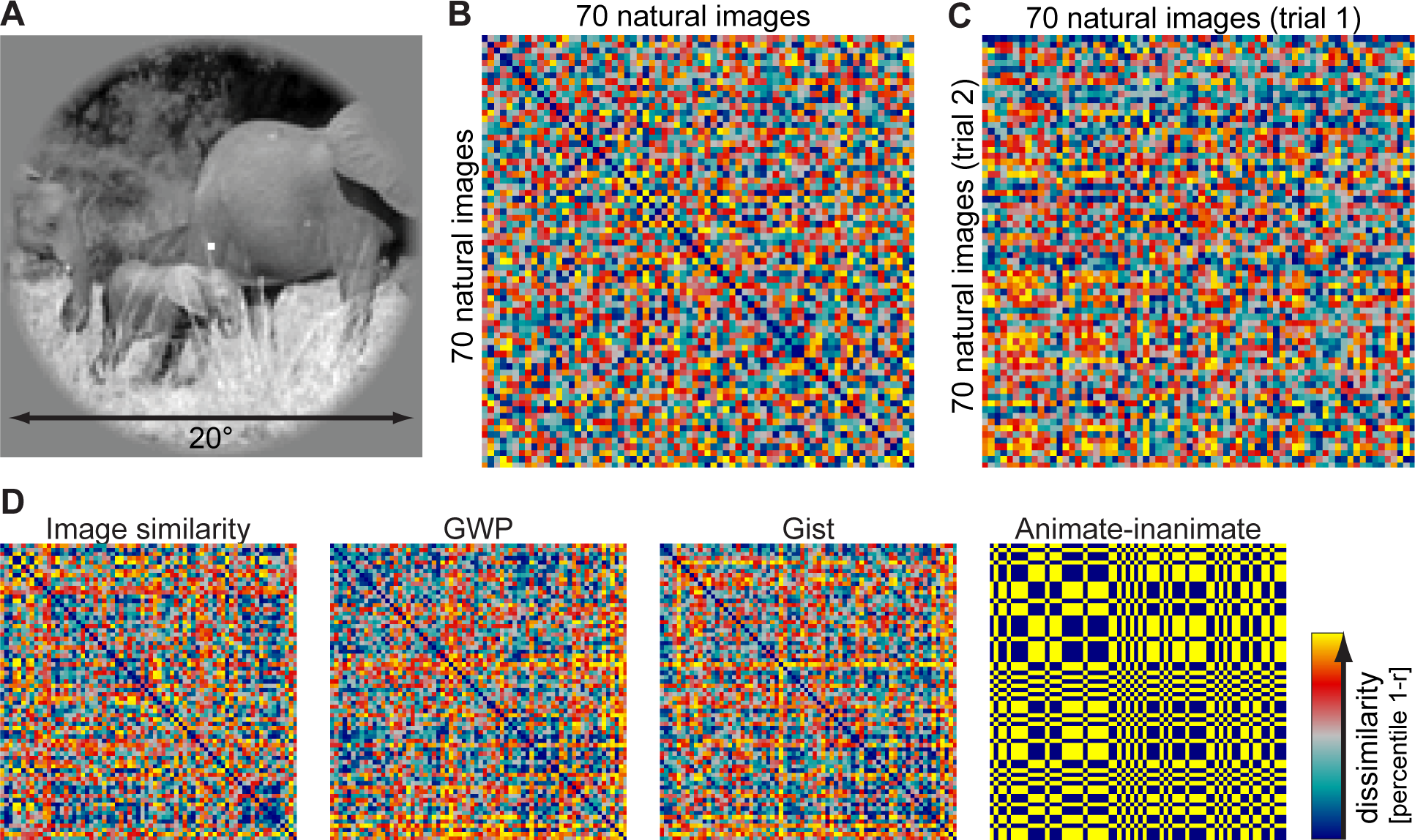
Representational similarity analysis and computational model predictions of natural image representations. A) An example stimulus image is shown. B) An example representational similarity matrix (RDM) is shown for visual area V1 of subject S1. The RDM captures the pair-wise dissimilarities between the response patterns elicited by the stimuli, here 70 different natural images. By definition, the RDM is symmetric and has a zero diagonal. C) In a split-data RDM, the dissimilarities are computed between separate presentations of the same set of stimuli. The diagonal of the split-data RDM reflects the replicability of the response patterns between the first and second stimulus presentation. D) Example RDMs for the four different models are shown for a set of 70 natural images (GWP = Gabor Wavelet Pyramid).

To study the ability to discriminate the natural images from the fMRI response patterns, we used split-data RDMs, where the pair-wise dissimilarities were computed between the two response patterns that were measured on different trials. Figure 3C shows an example of such RDM, where the diagonal reflects the dissimilarity of the response patterns between the first and second presentations of the same natural image. An index for the natural image discriminability was calculated as the subtraction of the mean of the diagonal values from the mean of the off-diagonal values. Exemplar discrimination index greater than zero indicates distinguishable response patterns for the natural images.

The replicability of the similarity structure captured by an RDM was assessed by comparing single-trial RDMs based on the two separate presentations of the same set of images. The RDMs were compared using Kendall’s tau-a rank correlation distances of the values in the upper (or equivalently the lower) triangle of the RDMs (for details on the different correlation-distance measures, please see (Nili *et al.* 2014)).

### Computational models

The visual area RDMs were compared to four different model predictions on the representational similarity structure: *image correlation similarity*, *Gabor wavelet pyramid model*, *Gist* and *animate–inanimate distinction*. In a model RDM, each cell reflects the dissimilarity of an image pair predicted by the computational model. Examples of the model RDMs are shown in Figure 3D. The comparison between a brain RDM and a model RDM was based on Kendall’s tau-a rank correlation distance of the values in the upper triangles of the RDMs.

#### Image correlation similarity

Each gray-scale image was converted to a vector and compared to each other image using correlation distance (1-correlation).

#### Gabor wavelet pyramid

The Gabor wavelet pyramid model was adopted from Kay et al. (Kay *et al.* 2008). Each image was represented by a set of Gabor wavelets of six spatial frequencies, eight orientations and two phases (quadrature pair) at a regular grid of positions over the image. To control gain differences across wavelets at different spatial scales, the gain of each wavelet was scaled such that the response of that wavelet to an optimal full-contrast sinusoidal grating is equal to 1. The response of each quadrature pair of wavelets was combined to reflect the contrast energy of that wavelet pair. The outputs of all wavelet pairs were concatenated to have a representational vector for each image. The pair-wise dissimilarities (1-correlation) of these vectors were computed to obtain the Gabor wavelet model RDMs for the natural images.

#### Gist

The spatial envelope or gist model aims to characterize the global similarity of natural scenes (Oliva and Torralba 2001). The gist descriptor is obtained by dividing the input image into 16 bins, and applying oriented Gabor filters in 8 orientations over different scales in each bin, and finally calculating the average filter energy in each bin. The gist descriptors for each natural image were compared to each other to obtain the Gist RDMs.

#### Animate–inanimate distinction

The natural images were labelled as animate if they contained one or several humans or animals, bodies of humans or animals, or human or animal faces. In the animate–inanimate model RDM, the dissimilarities are either 0 (identical responses) if both images are of the same category (animate or inanimate) or 1 (different responses) if one image is animate and the other is inanimate.

### Searchlight analysis

The main analyses were performed using pre-defined ROIs. To explore the effects more generally within the whole scanned brain volume, we performed searchlight analysis (Kriegeskorte et al. 2006). A spherical searchlight of 4.5-voxel radius was positioned at each location of the scanned brain volume. Within each location, the response-pattern-means and RDMs were extracted and correlated with the corresponding metrics from a reference ROI. Correlation maps were constructed from the results. This was repeated for the 25 experimental runs. The results were FDR corrected for multiple comparisons.

### Trial averaging

The effect of the number of fMRI response trials averaged was studied using a second set of fMRI data from the same subjects, where we had 13 trials for 120 natural images (for details, see the image identification data in (Kay *et al.* 2008)). The representational similarity analysis was applied separately for data from each experimental run with 12 different natural images. The trials were divided to two independent data sets (odd and even trials, the 13th trial excluded from the analysis), and the number of response patterns averaged was varied between 1 and 6. The averaged response patterns were used for representational similarity analysis.

## RESULTS

### Visual areas contain image information and exhibit replicable representational geometries

We studied fMRI response patterns for 1750 natural images in visual areas V1, V2, V3, V4, V3A, V3B, and LO. A portion of this data set has previously been analyzed with voxel-receptive-field modeling (Kay *et al.* 2008). The previous analysis showed that fMRI signals from the human visual cortex can be modeled by a Gabor wavelet pyramid. The distinct response patterns for the natural images were captured here by RDMs. Figure 3B shows, as an example, a V1 RDM in which each cell compares the V1 response-patterns elicited by two different natural images. An RDM captures the pairwise dissimilarities of the response patterns and can thus be directly compared to an RDM of a different brain region (without any need for voxel-to-voxel matching of the response patterns) or to an RDM describing the representational geometry of a computational model (Kriegeskorte, Mur and Bandettini 2008).

First to confirm the suitability of the data for representational similarity analysis, the distinctness of the response patterns for the natural images was studied using split-data RDMs in which each cell compares the response patterns between different trials for the same images (for an example, see Fig. 1C). The diagonal of the split-data RDM reflects the replicability of the response patterns for the repeated presentation of the same stimulus images. An exemplar-discriminability index was calculated by subtracting the mean of the diagonal values from the mean of the off-diagonal values. Figure 4A shows the results on exemplar discriminability. For all three subjects, the most distinct response patterns were found in V1. The exemplar discriminability indices were greater than zero in all studied visual areas (p < 0.05, one-sample t-test, for each visual area in each subject). This indicates that response patterns contain information about the stimuli.

**Figure 4.**
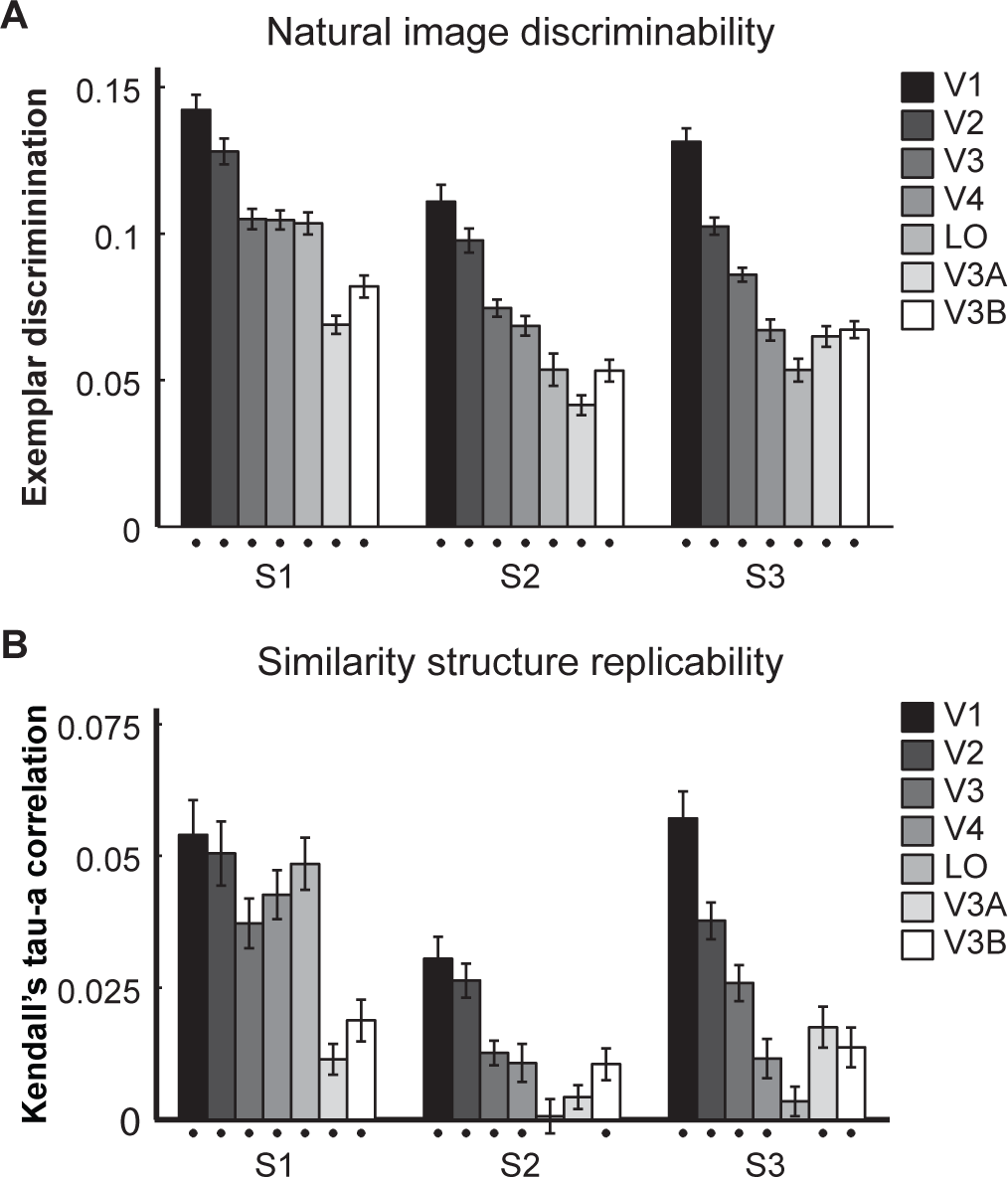
Distinct fMRI response patterns and replicable similarity structures for natural image stimuli. A) Results on the distinctiveness of the response patterns for the natural images are shown separately for the seven visual areas in each of the three subjects (S1, S2, S3). The error-bars indicate SEMs across the 25 experimental runs. The black dots below the bars indicate statistically significant results (t-test, p<0.05). B) Results on the replicability of the representational similarity structure for the natural image stimuli are shown separately for the visual areas and subjects. The error-bars indicate SEMs across the 25 experimental runs. The black dots below the bars indicate statistically significant results (t-test, p<0.05).

Beyond the mere presence of information about the image presented, we asked whether the RDMs were replicable. RDM replicability was assessed by rank correlation. RDM replicability would indicate that pairs of images are not all equally distinctly represented, but that some pairs are reliably represented as more similar than others. Figure 4B shows results on the replicability of the response-pattern dissimilarity structure. Here we compared single-trial RDMs based on two separate presentations of the stimuli. In all subjects, the RDM was best replicated in V1. For subject S1, the RDMs showed replicable structure in all studied visual areas (p < 0.05, one-sample t-test, n = 25). For subject S2, the single-trial RDMs did not show replicable structure in areas LO and V3A (p > 0.05). For subject S3, the RDMs did not show replicable structure in LO (p > 0.05) but the results for all other studied areas were significant (p < 0.05).

Taken together, these results confirmed that this data set with its rich sample of stimuli is suitable for representational similarity analysis.

### Gabor model explains early visual representation, LO exhibits categorical clusters

We compared the visual-area RDMs to RDMs constructed based on four different models: *image correlation similarity*, *Gabor wavelet pyramid model (GWP)*, *Gist* and categorical *animate–inanimate distinction*. Examples of the model RDMs are shown in Figure 1D. The image correlation distance was chosen as the simplest model that could explain variance in the response patterns elicited by the natural images. The Gabor wavelet pyramid is considered as the standard model of visual area V1 (Carandini et al. 2005). The Gist model has been suggested to capture the global features of natural scenes relevant for scene categorization (Oliva and Torralba 2001). The categorical animate–inanimate model was chosen based on previous studies suggesting this as a fundamental organizing principle of higher-level object responsive areas (Kiani et al. 2007; Kriegeskorte, Mur, Ruff, et al. 2008; Naselaris et al. 2012).

Figure 5 shows the results on the model fits to the empirical RDMs of different visual areas. The model fit was assessed using Kendall’s tau-a rank correlation between a visual-area RDM and a model RDM. This measure reflects how well the ranking of the response-pattern dissimilarities for a visual area was explained by the ranking of the dissimilarities of the model’s response patterns. In contrast to the voxel-receptive-field modeling approach (Kay *et al.* 2008), where each voxel response is predicted as a linear combination of model responses, the RSA approach obviates the need to estimate any parameters from the data here. In all subjects (Fig. 5), the V1 RDM was best explained by the Gabor wavelet pyramid model. The Gist model also captured some of the response variance but was not as good fit as the GWP. The GWP was also the best model for the response profiles of visual areas V2 and V3. In V4, all models, except the simple image correlation model, produced comparable model fits. The animate–inanimate model was the best-fitting model for area LO, especially for subject S1 (Fig. 5A). For subjects S2 and S3, the results on the RDM replicability (Fig. 4B) already suggested noisy LO data.

**Figure 5.**
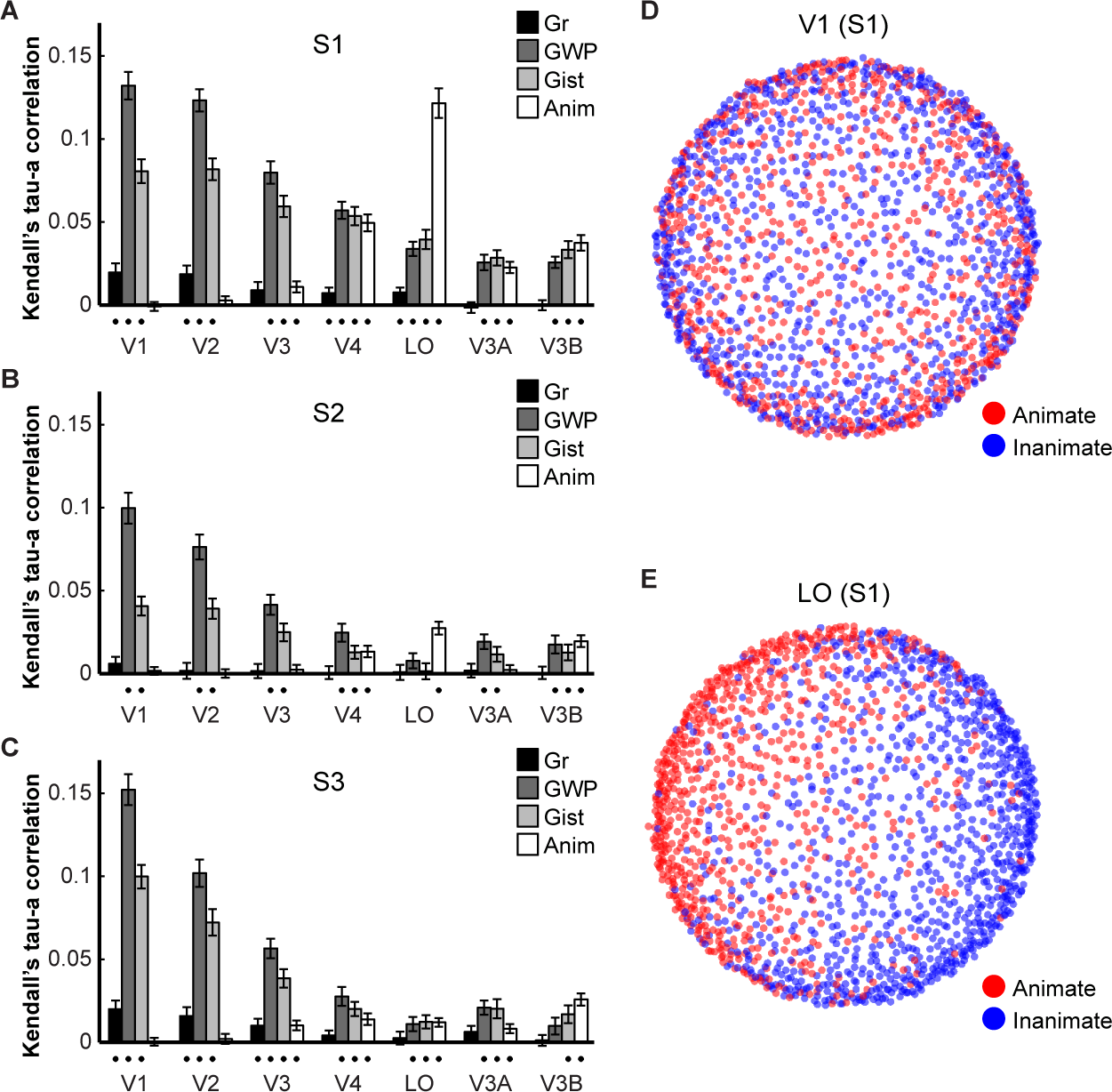
Relating computational models to cortical representations. A-C) Results on the comparisons between computational model predictions on the response pattern dissimilarities and the empirical results of different visual areas are shown separately for the three subjects. Each bar indicates the mean rank correlation between a model RDM (Gr = image correlation similarity, GWP = Gabor wavelet pyramid, Gist = spatial envelope model, Anim = categorical animate–inanimate distinction) and a brain RDM. The error-bars indicate SEMs across the 25 experimental runs. The black dots below the bars indicate statistically significant results (t-test, p<0.05). D-E) A multidimensional-scaling arrangement reflects the response-pattern dissimilarities in V1 and LO for the 1750 natural images (dissimilarity: 1-Pearson’s linear correlation, criterion: metric stress) labelled as animate (red) or inanimate (blue). A clear categorical clustering is evident in LO (E), but not in V1 (D).

Visualizations of the response-pattern dissimilarities for all 1750 natural images for visual areas V1 and LO from subject S1 are shown in Figures 5D-E. The stimuli were color-coded based on the animate–inanimate distinction. Each dot represents an individual stimulus. The distances between the dots reflect response-pattern dissimilarities. This multidimensional scaling visualization of the response-pattern dissimilarities is unsupervised (*i.e.*, without any assumptions of a categorical structure), and hence any observed distinctions are data-driven. The results show no categorical clustering of the animate and inanimate stimuli in V1. In contrast, a global grouping of the stimuli reflecting the categorical clustering between animate (red dots) and inanimate (blue dots) natural images is evident in area LO. This is consistent with a previous RSA study using 92 images of isolated objects (Kriegeskorte, Mur, Ruff, *et al.* 2008) and extends that finding by using a much larger stimulus set with the animate and inanimate objects embedded in natural scenes. Our results also show that the animate/inanimate distinction emerges at a later processing stage and is not due to low-level visual-similarity effect between the categories that would be present already at the level of V1.

The shift from the GWP to the categorical animate–inanimate distinction appears a plausible characterization of the changes in the representations of natural images across hierarchy of visual areas. Next we address how much of the response variance the selected models explain. Do the best-fitting models explain all replicable variance in the data, or is a replicable component left unexplained? This leads us to the question of the relative contributions of the intrinsic response fluctuations and stimulus-driven effects on response-pattern similarity.

### RDMs are highly similar among visual areas when estimated from the same trials

The transformation of representational similarity structure along the ventral visual stream can be investigated by comparing RDMs between visual areas. Figure 6 shows second-order dissimilarity matrices, where the RDMs of visual areas were compared to each other both within and between subjects, and to the two best-fitting model RDMs (GWP and animate–inanimate). In Figure 6A the visual area comparisons were done between RDMs constructed from response patterns that were estimated on the basis of the same (first) trials. Figure 6B shows the corresponding multidimensional scaling (MDS) arrangement where the RDMs of different subjects and of the two models are shown with different colours (S1 = blue, S2 = green, S3 = purple, Gabor wavelet pyramid model = black, categorical animate–inanimate model = red). The structure in the RDM as well as the MDS visualization show grouping of the visual areas based on the subject, not visual hierarchy. That is, the visual area RDMs were always most similar to the RDMs of other visual areas of the same subject. This would suggest that the models do not explain the stimulus representations in the visual areas very well. However, the following analyses show that the within-subject similarity of the RDMs was driven by the intrinsic dynamics (which are shared among areas within, but not between, subjects).

**Figure 6.**
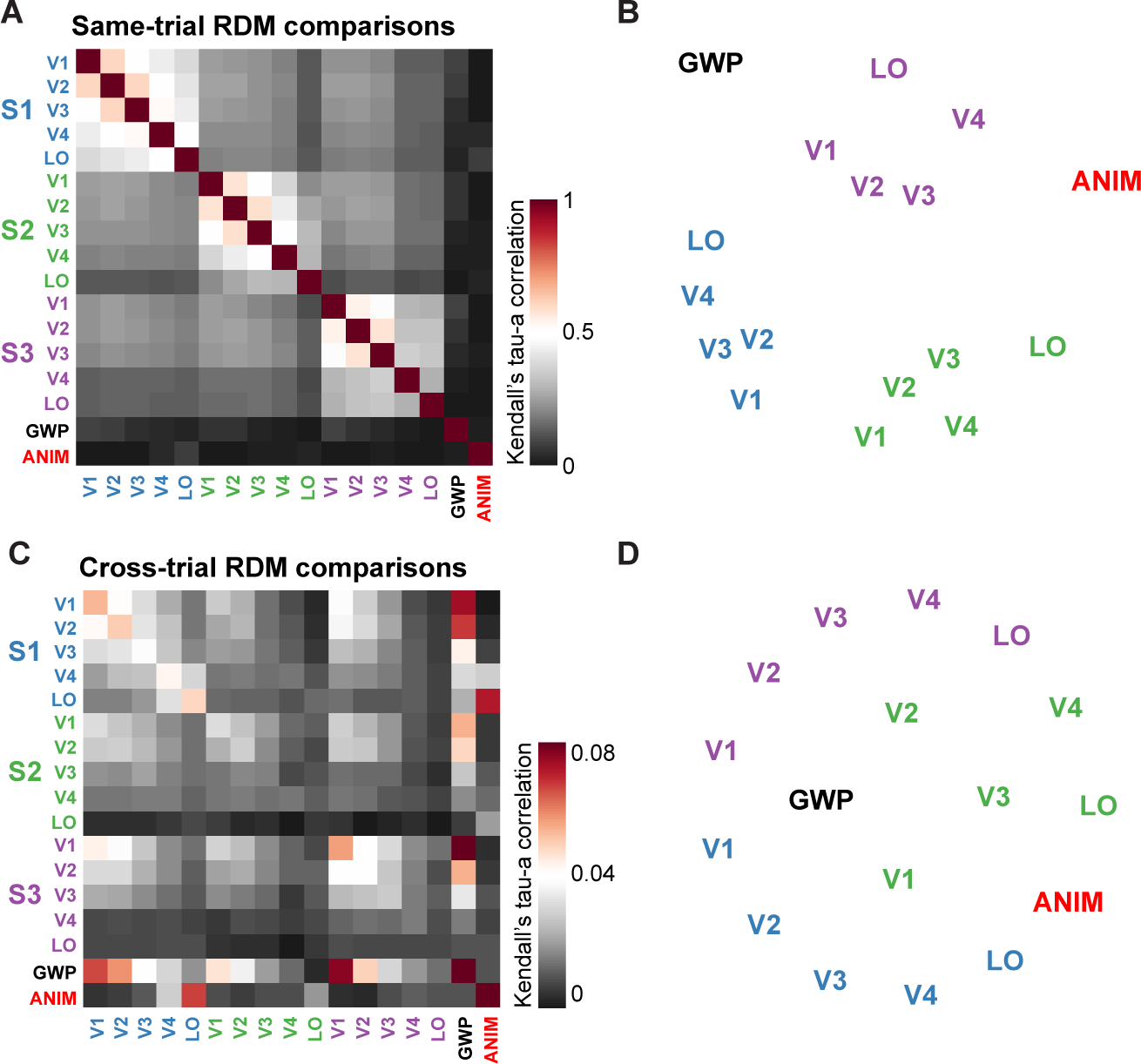
Relating natural image representations between different visual areas, subjects and model predictions. A) A second-order similarity matrix of RDMs of visual areas (V1, V2, V3, V4, LO) in all three subjects (S1, S2, S3) and the two best-fitting models (GWP = Gabor wavelet pyramid, Anim = categorical animate– inanimate model) is shown, and B) the corresponding multidimensional-scaling arrangement (metric-stress) of the representational dissimilarities. The distances reflect the representational distance between the representations. The visual areas in the three subjects are color-coded in different colors. C) A second-order similarity matrix of RDMs, where the effects of coherent trial-to-trial fluctuations were removed by comparing RDMs from separate trials, and D) the corresponding multidimensional-scaling visualization (metric-stress) of the representational relationships. Note that when the comparison was made between the visual-area RMDs constructed form the same trials (sharing intrinsic dynamics; A–B), the representations were most similar between the visual areas within the same subject. Whereas, when the comparison was made between visual-area RDMs constructed from separate trials (sharing only stimulus-driven effects; C–D), the V1 representations of all subjects, for example, were more similar to the GWP model than to the representations in the higher-level visual areas.

### RDMs are distinct among visual areas when estimated from separate trials

Next we compared the visual area RDMs constructed from response patterns that were estimated on the basis of separate trials. That is, the compared RDMs only shared the stimulus-driven component. Figure 6C shows the second-order dissimilarity matrix and Figure 6D the corresponding MDS arrangement of the RDM relationships. The results are very different from the same-trial results shown in Figures 6A–B. The use of separate trials to compare the RDMs revealed clearer distinctions between the visual areas and broke the grouping based on individual subjects. This shows that the clustering of the RDMs by subject in the previous analysis (Fig. 6B) did not result from each subject having an idiosyncratic stimulus-driven representation that is similar across his or her visual areas. Instead intrinsic dynamics unrelated to the stimuli (which are not shared between repeated presentations of the same stimulus or between subjects) account for the RDM clusters.

When the effect of intrinsic dynamics is controlled for by comparing only RDM estimates based on separate sets of trials between cortical areas, the transformation of the stimulus-driven representations can be accurately assessed. Figure 6D shows a multidimensional scaling arrangement that reveals the relationships among areas and models. RDMs now cluster by visual area, instead of by subject. The V1 RDMs of all subjects cluster around the GWP-model RDM. The LO RDMs of all subjects cluster around the animate–inanimate model RDM. With the similarities among visual-area RDMs no longer inflated by intrinsic dynamics, the arrangement reflects the stimulus-driven representations and accurately depicts the relationships among visual areas and models.

### The Gabor model explains the stimulus-driven component of the V1 representation

Figure 7 shows results of quantitative analyses of how well the V1 RDM was explained by the GWP model, by the V2 of the same subject, and by the V1 of the other subjects. The within-subject replicability of the V1 RDM (cf. Fig. 4B) provides a noise ceiling, which reflects the amount of explainable RDM variance. When the RDMs were estimated on the basis of the same trials (Fig. 7A), the response dissimilarity structure of area V2 (same subject) explained the V1 RDM results far better than the GWP model or the V1 results from other subjects (same trial here implies trials with the same stimulus presentation order; we will come back to this later). However, the low correlation between the V1 RDM with its replication suggests that most of the similarity between the V1 and V2 RDMs in the same subject was not driven by similar stimulus representations, but by the intrinsic dynamics and possibly other artefactual factors shared by the RDMs when same trials were used for estimating the RDMs.

**Figure 7.**
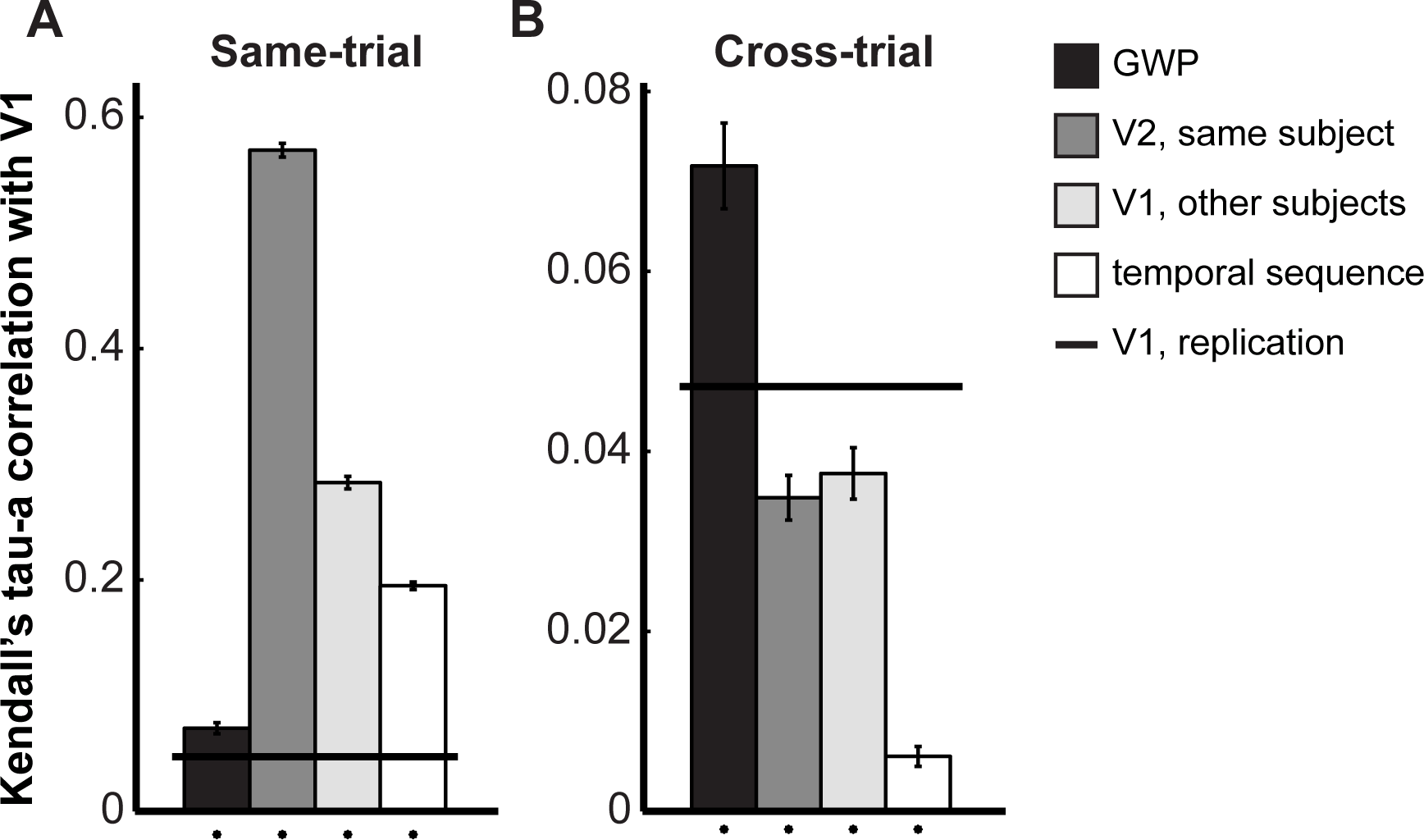
Same-trial RDM similarity is mostly driven by effects unrelated to the stimuli. A) Results are shown for same-trial RDM comparisons. The black bar shows the mean rank-correlation distance between single-trial V1 RDM and GWP model. The black line shows results on V1 RDM replicability, that is, the correlation between V1 RDMs constructed from separate trials and thus the stimulus-driven effects. The second bar shows the mean rank correlation between V1 RDM and V2 RDM of the same subject estimated from the same single-trial responses. The third bar shows the mean rank correlation of V1 RDMs from different subjects (“same trials”, denotes here the same stimulus presentation sequence). The fourth bar shows the mean rank correlation between V1 RDM and V1 RDMs of other experimental runs where different stimulus images were shown in the same temporal sequence. The last column thus reflects the contribution of the stimulus temporal sequence effects on the response-pattern similarity structures. The error-bars indicate SEMs across the 25 experimental runs. The black dots below the bars indicate statistically significant results (t-test, p<0.05). B) Results are shown for cross-trial RDM comparisons, that is, RDMs constructed based on separate trials (or different stimulus presentation order for the last two bars). Results on the GWP model (black bar) fit and V1-replicability (black line) are the same as in (A), note the different y-axis.

Figure 7B shows results from the cross-trial comparisons, that is, the comparisons of RDMs constructed on the basis of separate trials. The GWP model bar is the same as in Figure 7A (note the change of the vertical axis scaling). When the same-trial effects were removed, the V1 RDM was best explained by the GWP model. The same-subject V2 RDM and the between-subject V1 RDM (separate trial here implies trials with different stimulus presentation order; we will come back to this later) results are now below the ceiling-level as defined by the V1 replication (solid black line; same as in Fig. 7A). The GWP model outperformed the replication of the V1 RDM, reflecting the fact that both estimates of the V1 RDMs are noisy, whereas the GWP model is noise-free.

### Coherent response fluctuations within visual cortex

Correlated fMRI response fluctuations in the absence sensory stimulation are assumed to reflect intrinsic activity fluctuations within connected brain regions. We performed a searchlight analysis to explore the extent and specificity of the coherent response-pattern fluctuations and RDM correlations in our data. A spherical searchlight was positioned at each location of the scanned brain volume (Kriegeskorte *et al.* 2006). Within each location, the trial-to-trial response-pattern-mean as well as the RDM were extracted and correlated with the corresponding metrics from a reference ROI. The right LO was selected as the reference ROI, *i.e.*, the “seed region”. High correlation of response fluctuations were expected within visual cortex and especially with the corresponding region in the left hemisphere. In addition, left and right LO were expected to show similar stimulus-driven representations as reflected in similar RDMs estimated on the basis of separate trials. Similar visual representations cannot be assumed for low-level visual areas, where left and right visual fields are represented in opposite hemispheres.

Figure 8A shows the results on trial-to-trial response-pattern-mean correlations (Pearson’s linear correlation). The upper panel is equivalent to a functional connectivity analysis, where the seed “time course” is the trial-to-trial pattern mean from the right LO and this is correlated with trial-to-trial pattern mean of a spherical searchlight at each location. The correlations were high especially within the visual cortex. When the analysis was done between trial-to-trial mean signals for the same stimuli from different trials (lower panel in Figure 8A), the correlations were significant only around the location of the reference ROI. This suggests that there is a small stimulus-driven component also in response-pattern-means, but most of the same-trial response-mean correlations are not stimulus-driven. This finding is consistent with the presence of coherent intrinsic response fluctuations within the visual cortex.

**Figure 8.**
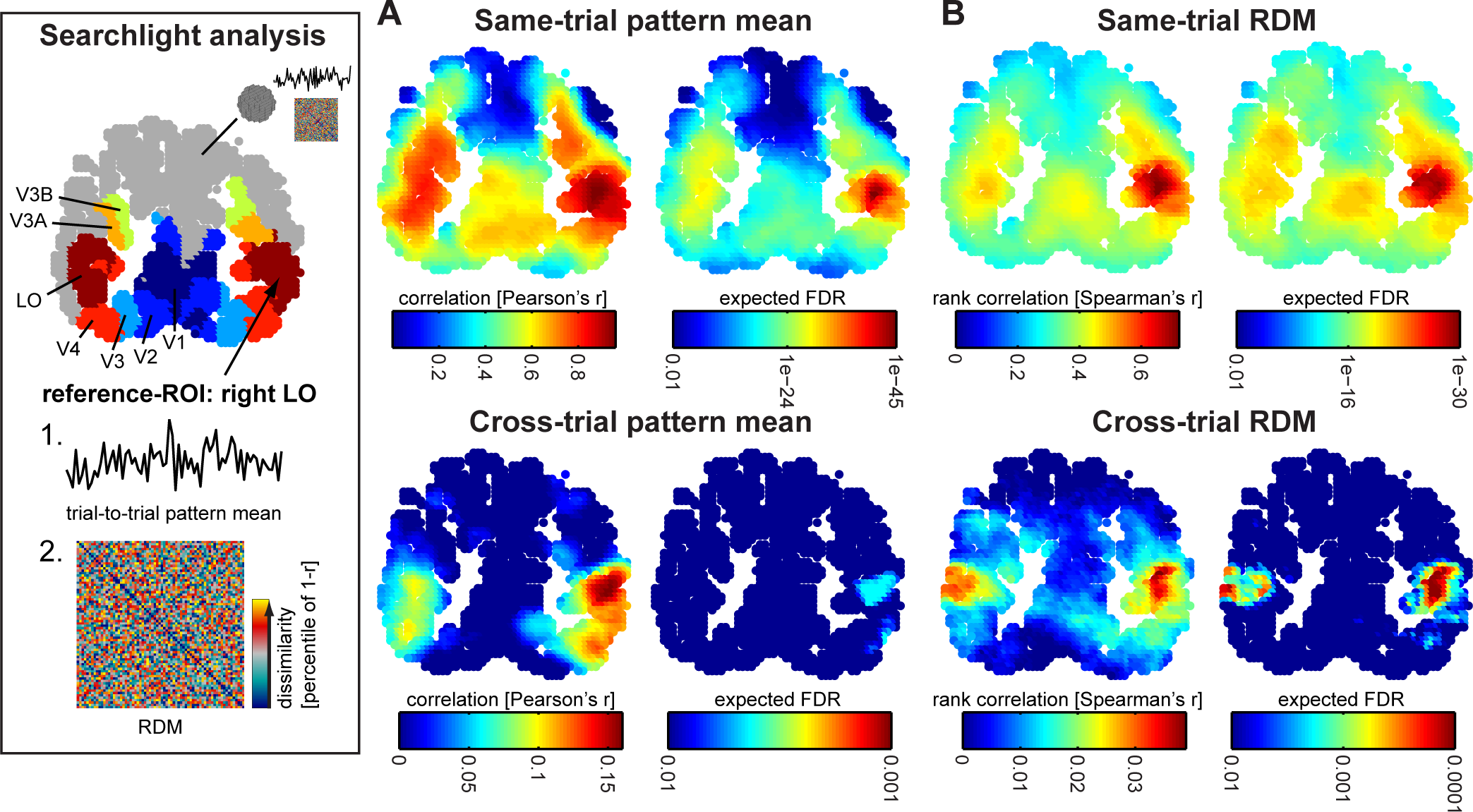
Searchlight analysis of the response-pattern fluctuations and RDM correlations across the visual cortex. A) The trial-to-trial response-pattern-mean signals from the right LO (subject S1) were correlated with trial-to-trial response-pattern-mean signals within a spherical searchlight at each location. The expected false-discovery rate maps show the significance of the correlations as evaluated from the 25 experimental runs and FDR corrected for multiple comparisons. The upper row shows the results for same-trial pattern mean correlations and the bottom row for different-trial pattern-mean correlations. Note the widespread response-pattern fluctuations in the same-trial responses across the visual cortex, and especially between the corresponding regions in the two hemispheres. B) The RDM of the right LO was correlated with RDMs within a spherical searchlight at each location. The upper row shows the results for same-trial RDM correlations and the bottom row for different-trial RDM correlations. When the reference-RDM was constructed from separate trials (bottom row), the searchlight analysis identified similar representations only in corresponding regions in the two hemispheres. Note the similarity of the same-trial RDM and same-trial response-pattern fluctuations across the visual cortex (upper rows A–B), likely reflecting the contribution from intrinsic cortical dynamics.

Figure 8B shows the searchlight analysis for RDM similarity. The RDM of the right LO was correlated (Spearman’s rank correlation) with RDM of a spherical searchlight at each location. The results for the same-trial RDM correlations (upper panel) resemble the results on the same-trial response-pattern-mean correlations shown in the upper panel of Figure 8A. The lower panel of the Figure 8B shows the results when the reference RDM from the right LO was correlated with RDMs constructed from response patterns within the spherical searchlights estimated on the basis of different trials for the same stimuli. This searchlight analysis picked up the right LO and the corresponding region in the opposite hemisphere, suggesting similar stimulus representations in these regions. These results support the conclusion that intrinsic cortical dynamics inflate the similarity of visual area RDMs and that the stimulus-driven similarity between the representations in different areas can be revealed by studying separate trials for the same stimuli.

### More dissimilar response patterns for trials more separated in time

In addition to the coherent fluctuations of the overall activity within the visual cortex, are there other non-stimulus related factors affecting the RDM similarity? We did find that the temporal sequence of stimulus presentation also had an effect on RDM similarity. In this data, the stimulus images in all experimental runs were different but they were presented using the same sequence (including timing of null trials and timing of the repetitions of the same stimuli). The correlation between V1 RDMs estimated on the basis of response patterns from different experimental runs is shown in Figure 7A (white bar). The correlation cannot be explained by stimulus-driven representation in V1 as the stimulus images in all runs were different. Moreover, the contribution of stimulus presentation sequence on RDM similarity likely explains also the difference in Figures 7A and Figure 7B for the between-subject comparison of V1 RDM similarity (light gray bars). Next we look more closely to the temporal effects on response-pattern dissimilarities.

The appearance of an RDM depends on the chosen stimulus order. If an RDM is ordered to follow the presentation sequence of the stimuli, do the temporal effects on the response-pattern dissimilarity become visible? Figure 9A shows a V1 RDM where the two trials for the 70 natural images within an experimental run were treated as separate conditions (RDM dimensions: 140 × 140). Here the ordering of the conditions in the RDM follows the numbering of the stimulus images (1-70). The first half corresponds to the first presentations of the images, the second half to the second presentations. In Figure 9B, the condition labels were reordered based on the temporal sequence of the stimulus presentation. In Figure 9C, these reordered RDMs were averaged across experimental runs. Because different natural images were shown using the same temporal sequence in all runs, the result should resemble an RDM, where the response patterns are similar for repeated presentations of the stimuli and dissimilar for all other comparisons (Figure 9D). This is indeed observed (detail 1 in Figures 9C, 9D). However, the RDM also exhibits a prominent structure of low dissimilarities around the diagonal, which indicates that response patterns for stimuli presented close in time evoked more similar response patterns than stimuli presented further apart. The similarity of response patterns acquired close together in time is consistent with a slow drift in pattern space that could be related to head motion and/or drifts of the state of the scanner and the subject’s physiological state. A similar drift was also seen in RDMs from other visual areas (Supplementary Figure 1; and for more detailed analysis of the drifts, Supplementary Figure 2). These global pattern drifts most likely contribute to the high correlation between RDMs of neighbouring visual areas (Figures 6A, 7A-B) when same-trial RDMs are compared. That is, same-trial RDM comparisons between brain regions are confounded by both the correlated response fluctuations and the temporal sequence-related pattern dissimilarity structure. This further reinforces the importance of using independent trials when drawing conclusions from exploratory analysis of RDM similarity between brain regions.

**Figure 9.**
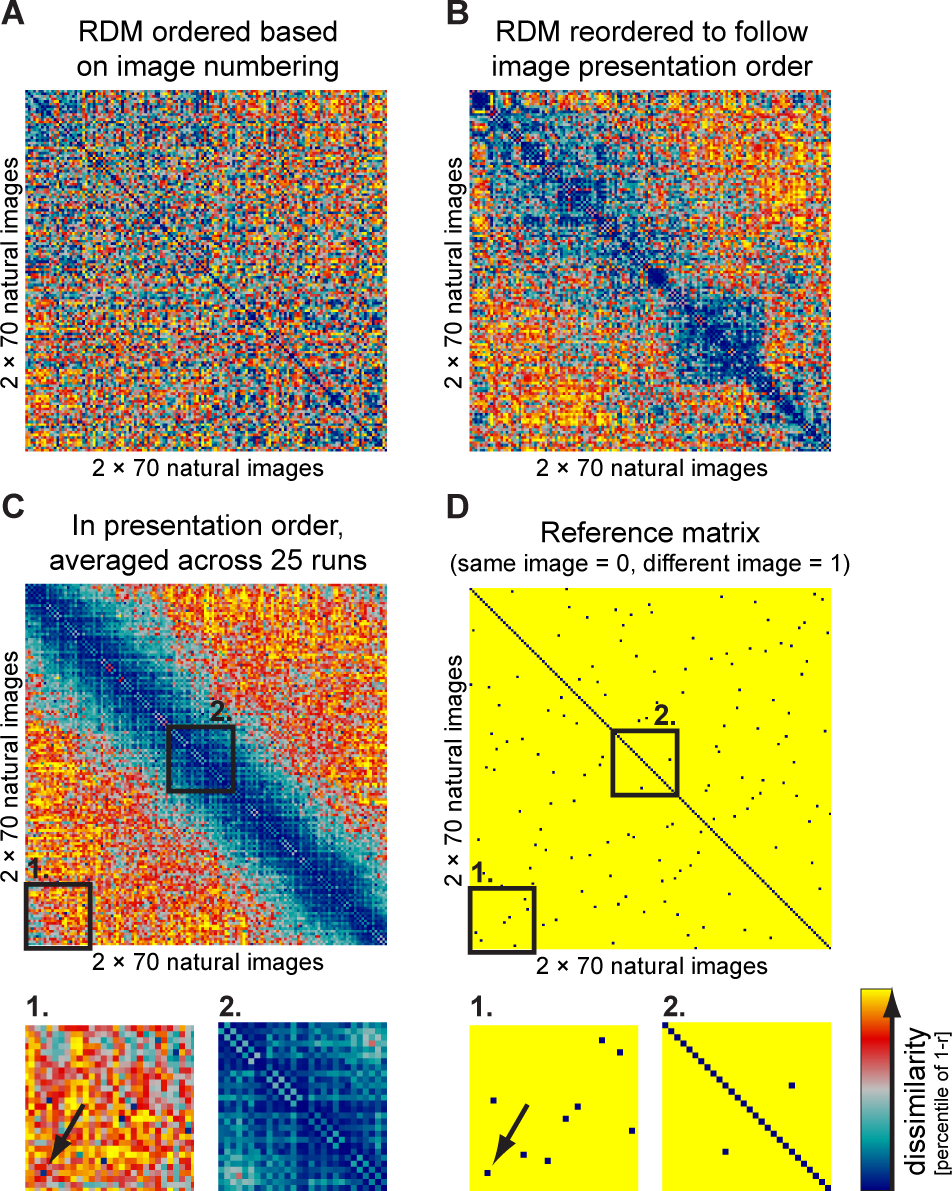
Trial-to-trial variability in response-patterns: more dissimilar response-patterns for trials more separated in time. A) A V1 RDM of subject S1 is shown for the first experimental run, where the two trials for each of the 70 stimuli were treated as separate conditions. The ordering of the condition labels in the RDM follows the original stimulus image numbering (1…70) with the second presentations of the same image set following the first presentation. B) The RDM shown in (A) is reordered to follow the temporal sequence of the presentation order of the 140 natural image stimuli. C) The reordered RDMs (as in B) were averaged across experimental runs. In each run, the stimuli were different, but the temporal sequence of the presentation order was same (see Supplementary Fig. 1 for other visual areas and subjects). The two black rectangles represent the two zoom-in regions. D) The reference RDM predicts identical response patterns for the repeated presentation of the same stimuli and different response patterns for other stimulus comparisons. The two black rectangles represent the two zoom-in regions. The arrows in the zoom-in regions point to one of the repeated presentations of the same stimulus (similar response patterns).

### Higher similarity of same-trial RDMs is not eliminated by averaging more trials

Thus far, our conclusions are based on single-trial RDM comparisons. Could the contribution of the coherent response fluctuations on RDM similarity between neighboring brain regions be eliminated by averaging more trials? We addressed this question with a data set where we had 12 responses for 120 natural images (Kay *et al.* 2008). The results are shown in Figure 10. The similarity of the same-trial RDMs was decreased when more trials were averaged (first column in Fig. 10). This is consistent with a contribution from correlated intrinsic fluctuations, which is expected to be reduced by averaging. At the same time, the similarity of RDMs estimated on the basis of separate trials was increased when more trials were averaged (second column in Fig. 10). Nevertheless, the correlation between the same-trial V1 and V2 RDMs remained much higher than the correlation between the RDMs estimated from separate trials. Hence, trial averaging appears insufficient to remove the non-stimulus-driven same-trial effects on RDM similarity.

**Figure 10.**
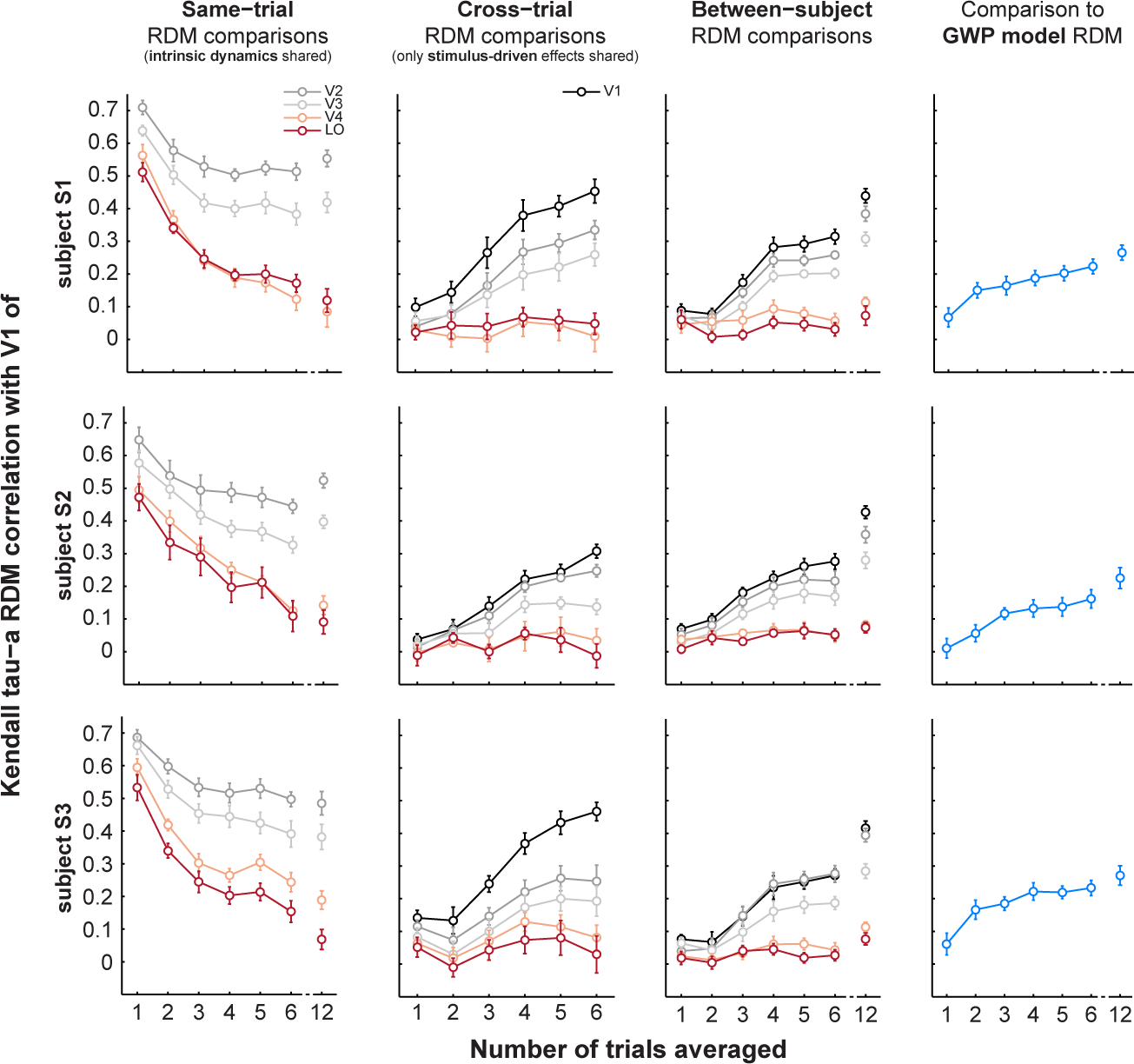
Trial averaging weakens non-stimulus-related effects and strengthens stimulus-related effects. Results on the representational similarity of V1 to other visual areas are shown for different numbers of trials averaged for the RDMs. Results for the three subjects are shown separately (three rows).The first column compares the V1 to other visual areas when the RDMs were constructed from the same trials (sharing intrinsic cortical dynamics). The x-axis shows the number of trials averaged for the RDMs. The second column compares the V1 to other visual areas when the RDMs were constructed from separate trials (only stimulus-driven effects shared among visual areas). Note the opposite effects of the trial averaging on the results shown in the first (same-trial) and second (cross-trial) columns. The third column compares the V1 representation in one subject to the representations in the other subjects (different temporal sequence for stimulus presentation; only stimulus driven effects shared). The last column compares the V1 representation to the representational similarity predicted by the GWP model.

### Representational dissimilarities in V1 are distinct from V2, and not fully explained by the Gabor model

Trial-averaging revealed a clear ordering of the RDM correlations (Fig. 10). The cross-trial RDM correlations are interpretable in terms of stimulus representational geometry. The V1 RDM was best explained by its replication in the same subject, followed by the same-subject V2 RDM and the other-subject V1 RDM. This suggests that the representational geometries are individually unique in V1. Finally, the basic GWP model’s representational geometry could not fully explain the V1 representation.

In summary, although trial averaging reduced non-stimulus-driven effects on the response patterns, it did not remove the effect of the same-trial coherent response fluctuations on the representational similarity between neighboring visual areas of the same subject. Using RDMs constructed from response patterns estimated on the basis of different trials, we were able to reveal subject-unique representational geometries, differences in representation between different visual areas, and the limits of the Gabor model for explaining V1.

## DISCUSSION

Intrinsic cortical dynamics are a major feature of cortical activity. Our results suggest five main conclusions: (1) Intrinsic dynamics exert a major influence on estimates of stimulus-related activity patterns and their dissimilarity structure. (2) The influence is such that representational dissimilarity matrices appear much more similar between two brain areas when estimated on the basis of the same trials than when estimated on the basis of a separate set of trials for each area. (3) A particular variant of intrinsic dynamics described in the literature, coherent fluctuations of activity across multiple areas, can account for this assimilation of the apparent representational geometries of two areas. Future studies will be needed to assess the intrinsic dimensionality of the intrinsic fluctuations and their functional role. (4) Response pattern estimates are also affected by substantial pattern drifts, which might be related to head motion, scanner state, or the subject’s physiological, emotional, or cognitive state. As a result of the pattern drift, two stimuli presented further apart in time will tend to be associated with more dissimilar pattern estimates. Single-trial-based RDM estimates therefore exhibit a stimulus-sequence-related component, which also creates spurious RDM correlations between different areas, when the same sequences have been used to estimate the RDMs. (5) The stimulus-driven component of the representation of a set of stimuli in two brain areas can be compared using a separate set of trials, which have been presented in independent random orders, to estimate the response patterns for each area. This approach avoids both the confound of correlated intrinsic fluctuations and the confound of sequence-related pattern similarity structure.

### Coherent response-pattern fluctuations between visual areas

From our data, we can only speculate the source of the coherent response-pattern fluctuations between the visual areas. A likely interpretation is the trial-to-trial variability in fMRI signal correlations that reflects underlying intrinsic spontaneous neural activity (Nir et al. 2008). The simulation and the observed coherent response-pattern fluctuations within the visual cortex support this conclusion.

We proposed that the contribution of the intrinsic fluctuations can be removed by comparing RDMs constructed of separate trials. Another approach would be to try to remove the widespread signal fluctuations from the data (Kay, Rokem, et al. 2013); this approach requires, however, multiple repetitions of the stimulus, and thus is not well-suited for the present data. Similarities between RDMs of the same visual area in different subjects can also reveal the amount of stimulus-related effects in the RDM. In between-subject RDM comparisons, each subject should have different random stimulus presentation order. This will prevent the temporal proximities of the stimuli from causing artefactual correlation of the pattern dissimilarities between the subjects.

We have emphasized that the intrinsic trial-to-trial variability in the response-pattern similarity can obscure the underlying stimulus-driven effects and thus confound results on representational similarities between brain regions. However, the trial-to-trial response fluctuations most likely have functional significance. Presumably the fluctuations are related to changes in the subject’s instantaneous state, such as attention and vigilance, and affect also the perception of the stimuli. Better reproducibility of fMRI response patterns has been previously associated with conscious perception (Schurger et al. 2010) and better memory (Xue et al. 2010) of visual stimuli. Furthermore, the effects of the temporal context of the stimuli on the response-pattern similarity could also be addressed in more detail in future studies. Visual cortex has been shown to show rapid adaptation effects to the structure of image stimuli at the single-neuron level (Muller et al. 1999) and at the level of neural populations (Benucci et al. 2013). This is likely also reflected in the pattern representations as measured with fMRI. Future work is needed to characterize the non-stimulus driven component of the response-patterns in more detail and to study its functional significance. The present experimental design of passive viewing of natural images is not suited to characterize for example the effects of attention (see, e.g., Ress et al. 2000) on the response-pattern similarity structure between visual areas.

### Relating fMRI results to computational model predictions of the underlying visual representations across stages of processing

RSA (Kriegeskorte, Mur and Bandettini 2008; Kriegeskorte 2009; Nili *et al.* 2014) and voxel-receptive-field modeling (Kay *et al.* 2008; Kay, Winawer, et al. 2013) are two complementary approaches to directly relate computational models with fMRI data. Voxel-receptive-field modeling aims to construct a computational model for each fMRI voxel and predict the responses for new stimuli, whereas RSA aims to predict the response-pattern similarity for a set of stimuli. We employed data that had been previously used as training data in voxel-receptive-field modeling. RSA does not require separate training data when the models have no free parameters to be estimated from the data. The model fit is determined by the correlation between the model RDM and a brain RDM. Here the stimulus set differed from previous RSA studies in that it was richer (>1000 images, as opposed to 96 in (Kriegeskorte, Mur, Ruff, *et al.* 2008); for a review, see (Kriegeskorte and Kievit 2013)). Our results confirm that this type of stimuli can be used with RSA to address questions on how representation of visual information is transformed along stages of the visual system and to test alternative computational models.

Our results on the model fits are consistent with previous studies showing that the representation of natural images in V1 can be explained by a Gabor wavelet model at the level of neural population codes (Weliky et al. 2003) and fMRI response patterns (Kay *et al.* 2008; Naselaris *et al.* 2009). The categorical clustering of responses for animate and inanimate objects in higher-level visual area is also consistent with previous work (Kiani *et al.* 2007; Kriegeskorte, Mur, Ruff, *et al.* 2008; Naselaris *et al.* 2012). Our results extend the previous studies by showing the shift in the best-fitting computational model across the hierarchy of visual areas as well as testing two other models (image correlation similarity and Gist) that did not perform better than the GWP or categorical animate– inanimate distinction in any studied visual area. Our results also confirm that low-level image similarity effects do not account for the categorical animate–inanimate distinction but this representation emerges in a higher-level visual area.

The present implementation of the GWP model is a simplification of what we already know about V1; the model does not include properties such as surround suppression (Cavanaugh et al. 2002) or cortical magnification (Duncan and Boynton 2003). However, taking into account the noise level in the data, it does a fairly good job at explaining the response dissimilarity in V1. However, when more trials were averaged for the RDMs, the limits of the GWP model in explaining the response variance became evident. Future studies should seek a computational account of both the prominent category divisions and the within-category representational geometry of LO. This might be achieved by testing a wider range of computational models (including newer models such as (Freeman et al. 2013; Kay, Winawer, *et al.* 2013)), with the aim also of characterizing the representational organizing principles of the intermediate-level visual areas (V2–4), in more detail. This would lead to a better understanding of the processing steps between the local, low-level image processing in V1 and the more global, category-selective representations in the higher-level visual areas.

## Conclusion

We found that coherent fMRI response-pattern fluctuations between visual areas can dominate representational similarities over stimulus-driven effects. Hence we suggest that representational similarity of brain regions should be addressed using response patterns estimated on the basis of separate fMRI trials. Here this approach revealed clear distinctions between the regions. More generally, our findings indicate that intrinsic cortical dynamics may have a significant contribution to representations as studied using multi-voxel fMRI pattern analysis.

